# Tuberculosis treatment outcome: The case of women in Ethiopia and China, Ten-Years Retrospective Cohort study

**DOI:** 10.1101/676924

**Authors:** Gebremeskel Mirutse, Mingwang Fang, Alemayehu Bayray kahsay, Xiao Ma

## Abstract

Every year tuberculosis kills above half million women all over the world. Nonetheless, the difference in the size of deaths among countries was not compared. Hence, this study is aimed to compare the death size of two countries. Socio demographic and clinical data of women treated for all form of tuberculosis in the past ten years 2007-2016 were collected from total of eight hospitals and six treatment centers of Tigray and Zigong respectively. Then, collected data were entered into SPSS version 21 then we estimated the magnitude of TB, level of treatment success and assessed factors associated with the unsuccessful TB outcome. In the past ten years, a total of 5603(41.5%) and 4527 (24.5%) tuberculosis cases were observed in Tigray and Zigong respectively. Of those with treatment outcome record a total of 2602(92%) in Tigray and 3916(96.7%) in Zigong were successfully treated. Total of 170 (6%) cases in Tigray and 36(0.8%) cases in Zigong were dead. In Tigray cases like retreatment (aOR, 0.29; 95% CI: 0.16-0.53) and multi drug resistant (aOR, 0.31; 95% CI: 0.003, 0.27) were less likely to show treatment success. But, HIV co-infected TB cases (aOR, 3.58; 95% CI: 2.47, 5.18) were more likely to show treatment success. In Zigong, women with MDR TB (Adjusted OR, 0.90; 95%CI: 0.24, 0.34) were less likely to show treatment success. On the other hand women in the age category of 15-49 (adjusted OR, 1.55; 95% CI: 1.08, 2.206) showed treatment success. Big number of tuberculosis cases and death were observed in Tigray comparing with Zigong. Hence, a relevant measure should be considered to improve treatment outcome of women in Tigray.

## Introduction

Despite the discovery of effective and affordable chemotherapy (1) tuberculosis kills 1.5 million people every year the death tall for women was 41.3% of the total death (2). The gender difference in tuberculosis infection was not well understood. Nevertheless, TB kills more women annually than all the causes of maternal mortality combined (2). In recent times, every year, at least 3.5 million women and children develop active TB among these 1.2 million cases died and more were left severely disabled (3-5).

Globally implementation of Direct observed treatment [DOTs] were saved 2.2 million of women and children (6) however there are enormous difference on the number of life saved and its factors affecting among regions. Few reports and studies attempts to compare and display the regional difference but the majorities were crude. For instance, WHO categorize Ethiopia and China among high TB burden countries (7-9) but its prevalence and treatment success reports were not specific for this group it tells about general population. Thus, the prevalence for Ethiopia was 192/100,000 and 67/100,000 was for China (10, 11) and the treatment success rate was 89% for Ethiopia and 94 % for China (10). In 2010 there was a regional report from high TB burden countries and out of 22 countries only 10 country report contain specific data about women and children that time China notified a total of 869 092 TB cases out of this 17% were women and 0.8% were children (5) at the same year Ethiopia notified a total of 150, 221 TB cases yet the size for women and children were not specified (5, 12).

Tigray which found in Northern part of Ethiopia (13) in 2015 notified a total 9,594 TB cases of both sex to a national TB control program. Accordingly 2,043 (21%) were smear positive with the cure rate of 74% (1,235) and 344 (4.2%) TB cases were died (14). Zigong which is located in southeastern Sichuan and which is home of large number of TB cases (15, 16) in 2015 notified 1738 TB cases and among this 399(22%) were smear positive with the cure rate of 385 (96%) and among total 22(1.2%) cases were died.

There are factors which identified as causes of unsuccessful treatment outcome for general population among these Retreatment cases, HIV co-infection, TB type and age were mentioned repeatedly however, this factors may not be found equally in all regions (17).

Generally the information in the above did not indicate the burden of TB in women specifically this implies that the existing study results and reports were crude. Thus, globally this time we lack specific proof which shows level of treatment outcomes and its factor affecting in these vulnerable group.

Declaring the above reasons, re-examining and comparing age and sex-aggregate data maintained by TB programs of these countries will be worth enough to look the profile, burden, treatment success and its factors affecting with in women. Moreover, finding of this study will help in tackling the limitation, shearing experience between countries and devise strategy to improve TB prevention and treatment program.

## Study settings and methods

### Tigray region

This study was conducted in Tigray (Ethiopia) and Zigong (China). Ethiopia is located in the Horn of Africa and is bordered by Kenya, Somalia, Sudan, Eritrea, and Djibouti. Administratively, Ethiopia is divided into nine regional states and two city administrative councils. The current population size was estimated 100 million(7, 18). Tigray is one of the nine national regional states of Ethiopia which is bordered by Eritrea in north and Sudan in the west. The region is administratively divided into seven Zones and 52 districts.

In 2010 among total population 2,441,158 (50.7%) were females with the total fertility rate of 5.1, agriculture is the main means of subsistence in the region in which 85% of the population lives in rural area.

In this region health care services are delivered through 1 specialized hospital, 15 general hospitals, 20 primary hospitals, 204 health centers, 712 health posts [village clinic] and 500 private health facilities. Then, in the region a total of 8,279 health professionals founds and 226[2.7%] were doctors (14). In Ethiopia Directly Observed Treatment (DOTS) was started in 1992 as a pilot and currently achieves 100 percent geographical coverage and recently 92% of public hospitals and health centers offer DOTS (19). The role of health facility in TB prevention and control was not centralized that means all hospitals and majority of health centers which have the diagnostic technology are allowed to diagnosis TB and providing DOTs service(17, 19, 20). Though, the role of health posts (village clinics) was limited only they provide health education, refer TB suspects for investigation and collect sputum smears, retrieve absentees/defaulters and in few place they can provide DOTs for case who is very far from health institution(21).

### Zigong region

China’s National TB control Program started to implement the international recommended directly observed treatment, short-course (DOTS) strategy in 1991, and expanded the DOTS program to the entire country by 2005 (8). Zigong is found in south-west China Sichuan province and currently this county has a total population 3.28 million. The existing health care system was organized into a three-tier health care delivery system and tuberculosis control program is centralized. Thus, the basic unit of TB health care is the specialized County TB dispensary (CTD) with the responsibilities of TB diagnosis, treatment and patient management guided by the National TB control program(11). Whereas, the non-CTD’s role in TB control program is to refer suspected TB patients to CTDs (22) all patient took anti TB drug three times per week (8) and DOTs observers get paid 60Yuan (US$1 ≈CNY7) per TB patient for the standardized treatment regimen of ≥6 months (23) The recommended treatment regimen for new TB case and retreated TB cases were the same 2HRZS/4HR and 2HRZSE/6HRE (16)

### Study design, sampling technique and data collection

Using retrospective cross-sectional study design we reviewed all form of TB cases of women and children (age greater than 15 years old) treated in the years of January 2007 to December 2016 in both countries. In Tigray DOTs is decentralized. So, all health facilities are allowed to provide DOTs but the data is not in digital form. Where as in Zigong DOTs services are not decentralized it is provided only in specific health facility but the data is available in digital form. Hence, considering the logistic constraint in Tigray among 16 hospitals eight hospitals that provide DOTs service for ten years and above were randomly selected then trained data collectors and supervisors were assigned to collect the information. Nevertheless, in Zigong there are only six TB treatment centers and all information about TB were found in CDC thus all data was extracted from Excel sheet.

### Data analysis

After checking the completeness, data was entered and analyzed using SPSS Version 21. Then, descriptive analysis such as frequency, mean and standard deviation were computed and compared. Additionally, binary logistic regression was executed to examine the association of independent variable with unsuccessful treatment outcomes. Hence, P-value less than of 0.05 was used as significant value. Finally, variables significant in binary logistic regression were analyzed again using multiple logistic regressions to identify variables which augment unsuccessful treatment outcomes in women and children.

### Ethical clearance

The study was passed through the ethical approval procedure of Sichuan University College of public health Chengdu, China and CHS/IRB of Mekelle University Ethiopia.

## Result

### 1. Socio demographic and clinical characteristic of women

The past ten years (January 2007-December2016) in Tigray a total of 13,435 and in Zigong 18,423 TB cases were identified. Among this 5603(41.7%) cases in Tigray and 4527(24.5%) in Zigong were women in the age category of 15-49. The mean age of case in Tigray was 36 ±15 years and in Zigong was 44 ±17years. Looking the age category in Tigray 4274 (76.3) and Zigong 2798(61.8) of women were in the age category of 15-49.

In addition among all TB cases 1175(21%) in Tigray and 2151(47.5%) in Zigong were pulmonary positive cases. Then, 21(0.4%) in Tigray and 16 (0.3%) in Zigong were MDR-TB. Also, 1048(18.7%) TB/HIV cases were identified in Tigray whereas no HIV documentations were found in Zigong. (table 1)

**Table 1.**
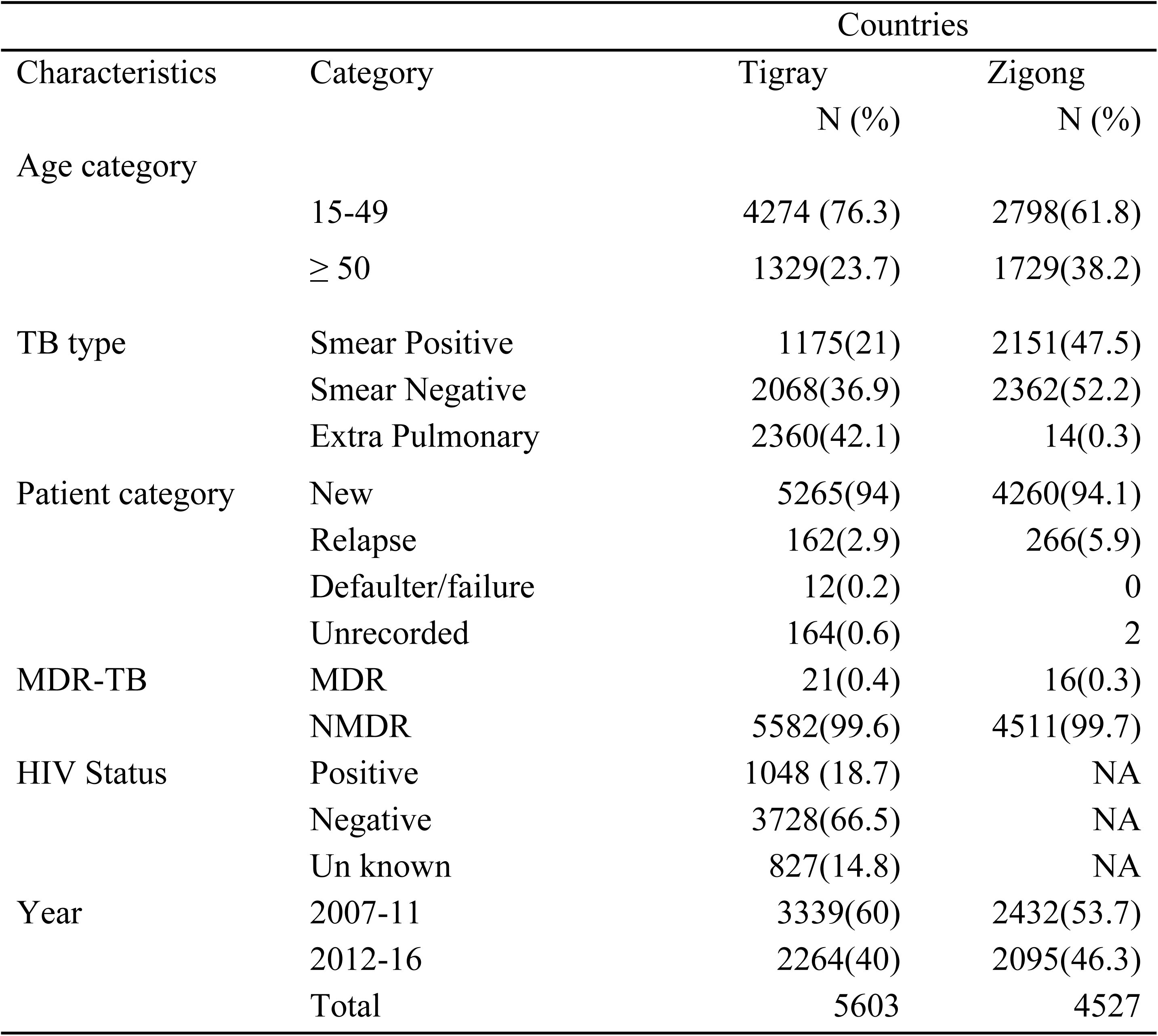
General characteristic of women treated for tuberculosis in Tigray and Zigong January 2007-December 2016[N=5603 and N=4527].

| Characteristics | Category | Countries |  |
| --- | --- | --- | --- |
|  |  | Tigray<br>N (%) | Zigong<br>N (%) |
| Age category |  |  |  |
|  | 15-49 | 4274 (76.3) | 2798(61.8) |
|  | ≥ 50 | 1329(23.7) | 1729(38.2) |
| TB type |  |  |  |
|  | Smear Positive | 1175(21) | 2151(47.5) |
|  | Smear Negative | 2068(36.9) | 2362(52.2) |
|  | Extra Pulmonary | 2360(42.1) | 14(0.3) |
| Patient category |  |  |  |
|  | New | 5265(94) | 4260(94.1) |
|  | Relapse | 162(2.9) | 266(5.9) |
|  | Defaulter/failure | 12(0.2) | 0 |
|  | Unrecorded | 164(0.6) | 2 |
| MDR-TB |  |  |  |
|  | MDR | 21(0.4) | 16(0.3) |
|  | NMDR | 5582(99.6) | 4511(99.7) |
| HIV Status |  |  |  |
|  | Positive | 1048 (18.7) | NA |
|  | Negative | 3728(66.5) | NA |
|  | Un known | 827(14.8) | NA |
| Year |  |  |  |
|  | 2007-11 | 3339(60) | 2432(53.7) |
|  | 2012-16 | 2264(40) | 2095(46.3) |
|  | Total | 5603 | 4527 |

### 3. Women clinical character and tuberculosis treatment in Tigray and Zigong

Over the study period, total of 5603(41.7%) and 4527(24.5%) tuberculosis (TB) cases were registered in Tigray and Zigong respectively. Then, in Tigray among all cases 2728(48.7%) TB cases were transferred to their nearby health facilities, 71(1.3%) have no record of their treatment outcome and in Zigong 480 (10.6%) case their treatment outcome were not recorded. Therefore, transfer out and cases with unknown treatment outcome were not included in the analysis. So, a total of 2804(50%) cases from Tigray and 4047(89%) from Zigong were involved in the analysis. Accordingly, 2602(92%) in Tigray and 3916(96.7%) case in Zigong were successfully treated. The cure rate of pulmonary positive cases out of 528 cases 477[90%] in Tigray where as in Zigong among 1891cases 1801 [95%] were cured. (Table: 2)

**Table 2.** women clinical factors and level of unsuccessful treatment outcome in Tigray and Zigong from January 2007-December 2016[N=2804 and N=4047].

| Characteristic | Country |  |  |  |  |  |
| --- | --- | --- | --- | --- | --- | --- |
|  | Total | Tigray<br>Unsuccessful<br>N (%) | Death | Total | Zigong<br>Unsuccessful<br>N (%) | Death |
| **Type TB** |  |  |  |  |  |  |
| P-Positive | 528 | 51(9.6) | 35(6.6) | 1889 | 60(3.2) | 22(1.2) |
| P-Negative | 1062 | 83(7.8) | 73(6.8) | 2156 | 70(3.2) | 13(0.6) |
| E- pulmonary | 1214 | 71(5.8) | 62(5.1) | 2 | 1(50) | 1(50) |
| **Treatment** |  |  |  |  |  |  |
| New | 2709 | 178(6.5) | 158(5.8) | 3822 | 122(1.7) | 32(0.83) |
| Re treatment | 95 | 24(25.2) | 12(12.6) | 225 | 9(4.0) | 4(1.7) |
| **Age** |  |  |  |  |  |  |
| 15-49 | 2155 | 156(7.2) | 134(6.1) | 2494 | 69(2.8) | 11(0.4) |
| ≥50 | 649 | 46(6.9) | 36(5.4) | 1553 | 62(4.1) | 25(1.6) |
| **Drug resistance** |  |  |  |  |  |  |
| MDR | 7 | 6(75) | 3(42.8) | 13 | 3(31.2) | 0 |
| NMDR | 2797 | 196(7.0) | 167(5.9) | 4034 | 128(3.2) | 36(0.9] |
| **HIV status** |  |  |  |  |  |  |
| Positive | 596 | 60(10) | 56(9.3) | 0 | NA | NA |
| Negative | 1838 | 88(4.7) | 70(3.8) | 4047 | NA | NA |
| Not tested | 370 | 54(14.5) | 49(13.2) | 0 | NA | NA |
| **Total** | **2804** | **202** | **170** | **4049** | **137** | **36** |

### 4. The trend of treatment success in women

The past ten years trend of treatment success was assessed the percentage of treatment success in Tigray was between 86%-98% and for Zigong was between 94%-98%. In Tigray the lowest treatment success was seen in the year 2007 which is 81% and in Zigong the lowest treatment success was seen in 2013 and its percentage was 94%. The overall treatment success in Tigray was 92% were as in Zigong it was 96.6%.

The graph for treatment success indicates in Tigray the past ten years there was constant increment then sharp decrease in the year 2016. But in Zigong it was a constant increment. (Fig: 1)

**Figure 1.**
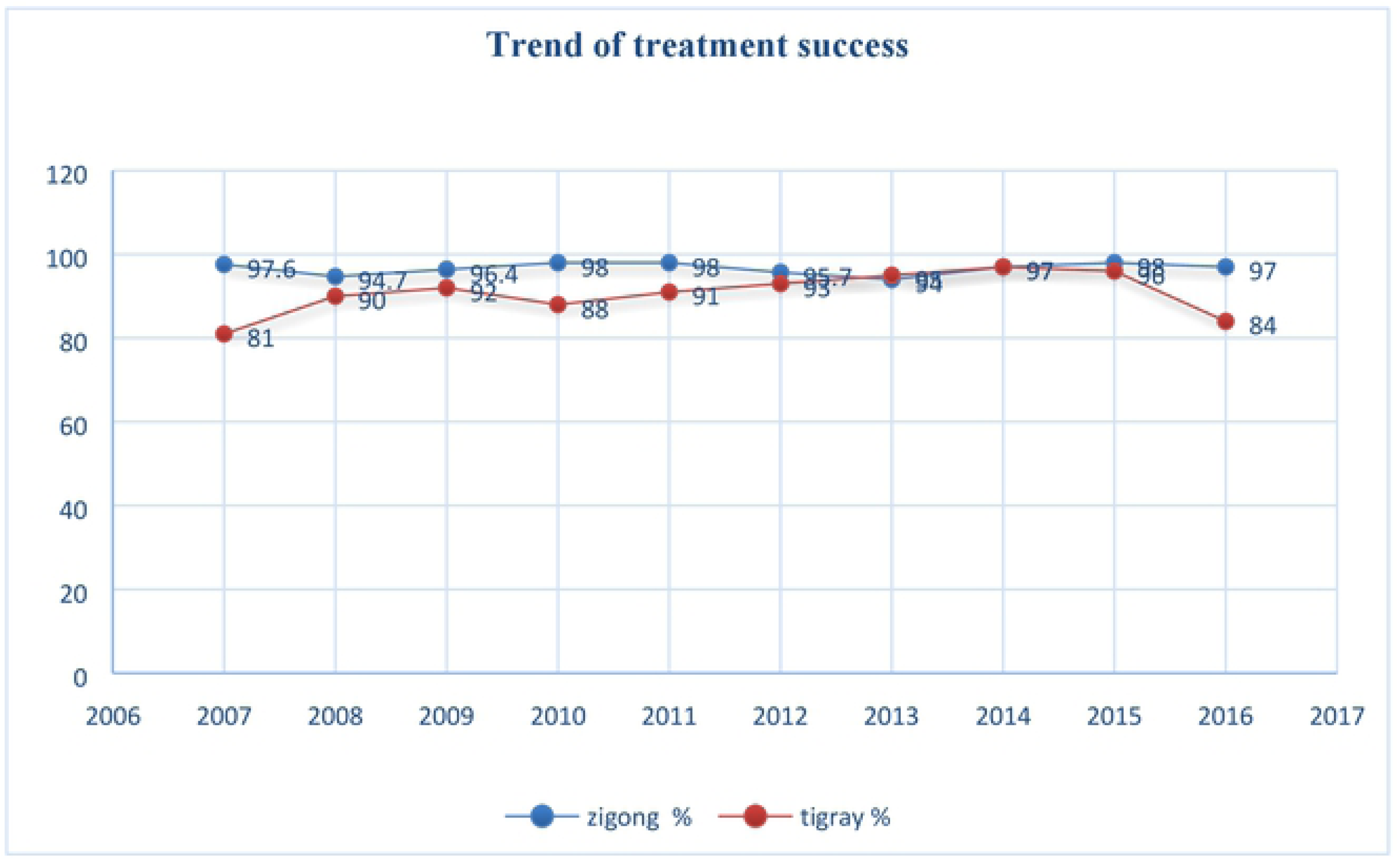
Trend of women treatment success in Tigray and Zigong January 2007-December 2016 N= 2084 and N=4047

### 5. Trend of tuberculosis death in Tigray and Zigong

In the past ten years a total of 170 [6%] case in Tigray and 36 [0.9%] cases in Zigong were reported died which is 6:1 ratio. Besides, in Tigray the peak death was seen in the year 2007 [12%] and in Zigong were in 2011 (0.57%). looking the trend of death in Zigong it was constant with the average death of 0.8 per year where as in Tigray it was a decreasing pattern and the average death per year was 5.8 per year of 100 cases. (Fig:2)

**Figure 2.**
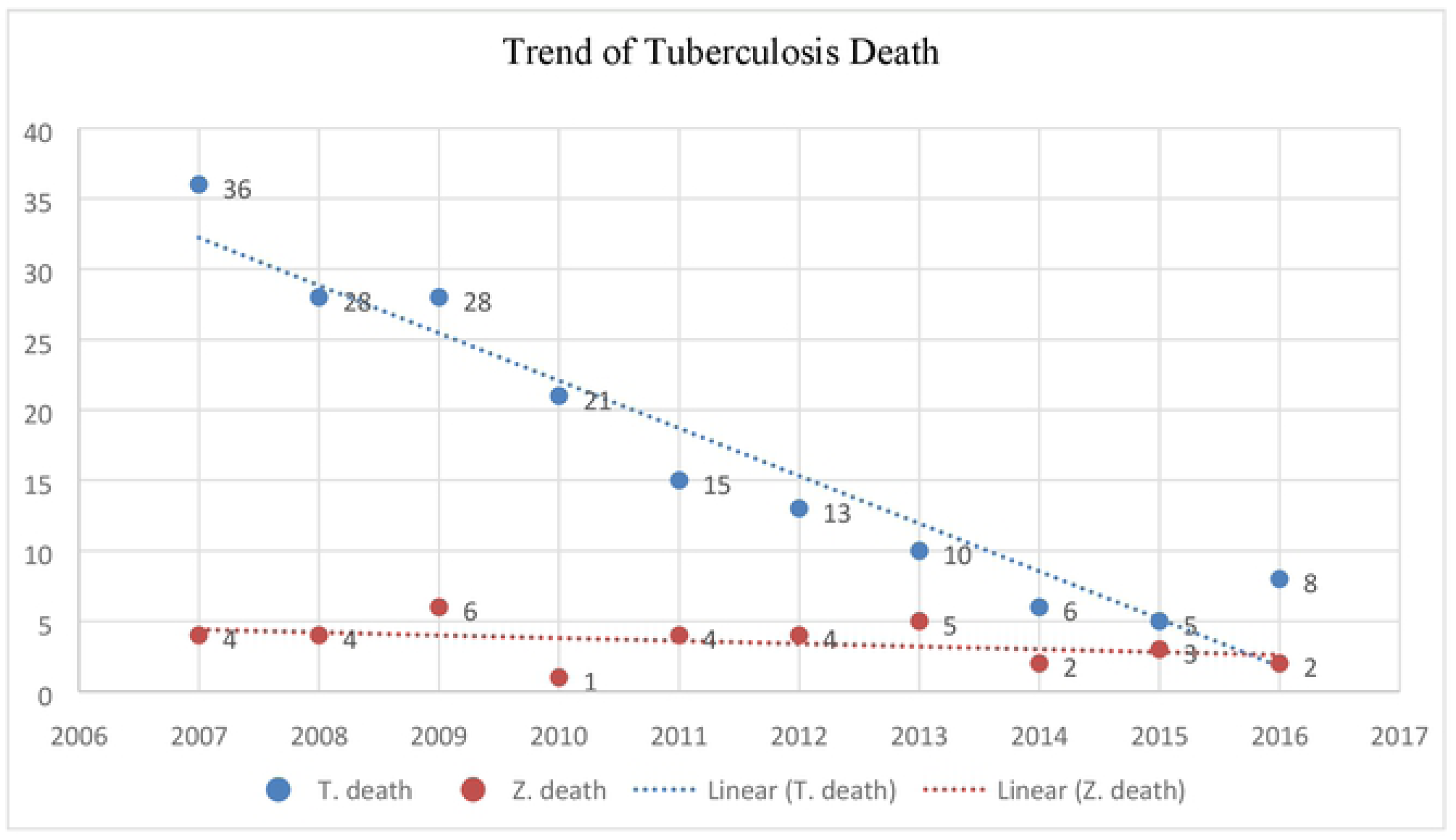
Trend of tuberculosis death in the past ten years (January 2007-December 2016) in women, Tigray N= 2084 and Zigong N= 4047

### 6. BLR: Sociodemographic and clinical Factors associated with unsuccessful treatment outcomes in women

In this study, we did bivariate logistic regression to identified factors that have association with unsuccessful treatment outcomes for both countries.

Accordingly, in Tigray those TB cases in retreatment category MDR cases, pulmonary positive and negative were more likely to show unsuccessful treatment outcome comparing with their counter parts. But variable labeled as HIV positive cases and HIV negative comparing with those unknown HIV status have less likely to have unsuccessful treatment outcome.

In Zigong MDR cases and age between 15-49 TB have more likely to have unsuccessful treatment outcome compared with others. (Table:3)

**Table 3.**
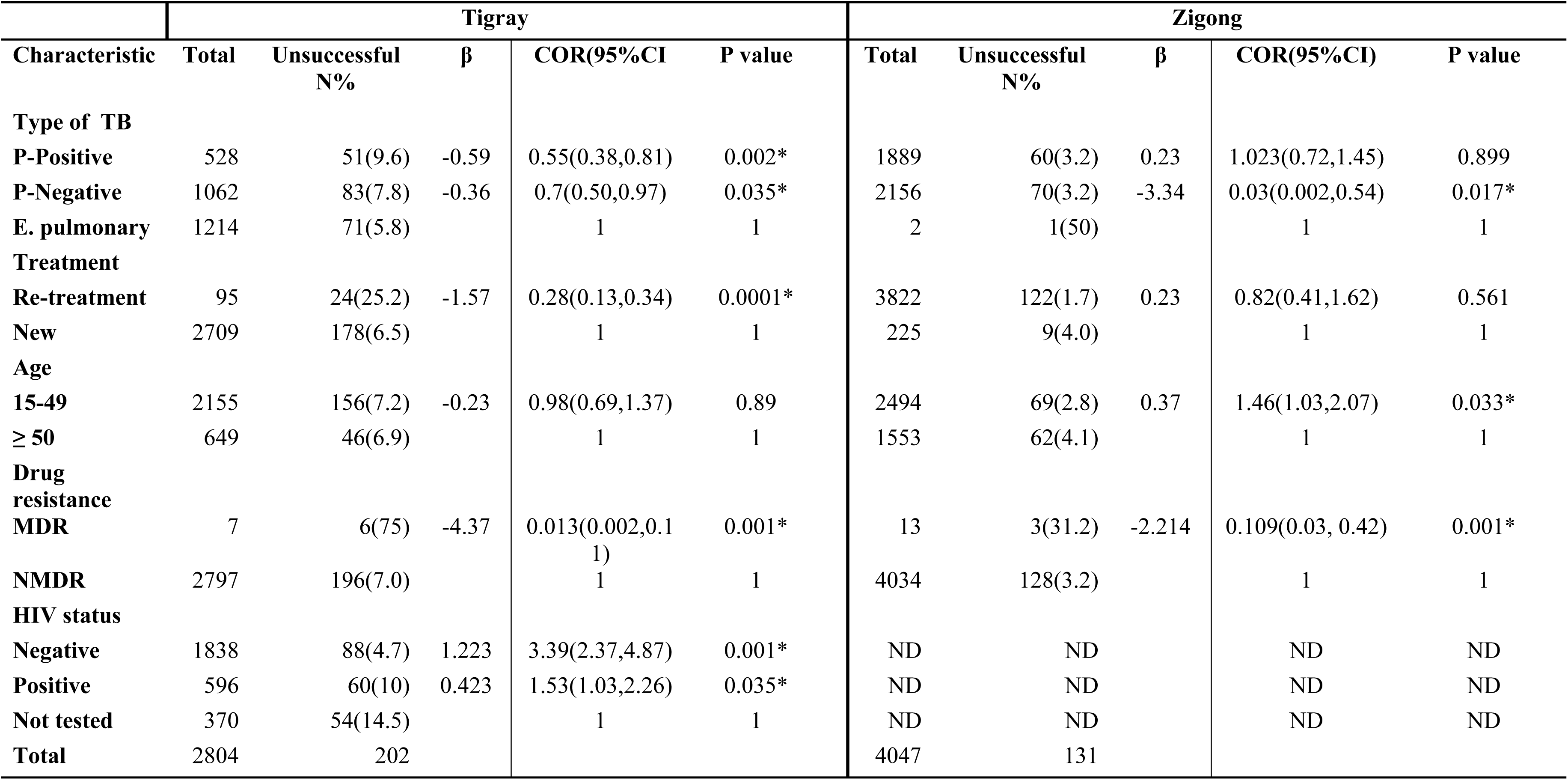
MLR Factors associated with treatment outcome of women in Tigray and Zigong January 2007 December 2016 N=2084 and N= 4047.

### 7. MLR: factors associated with treatment outcome of women in Tigray and Zigong

Factors significant at P-value < 0.05 in the bivariate logistic regression were took and analyzed again in multivariate regression to identify the predictor variables. Then, in Tigray treatment success was less likely for women who were categorized as retreatment (adjusted OR, 0.29; 95% CI: 0.16-0.53) compared to new cases, women with multi drug resistant (adjusted OR, 0.31; 95% CI: 0.003, 0.27) compared with non-drug resistant. But, HIV co infected TB cases were 3.58 times more likely to have treatment success (95% CI: 2.47, 5.18) compared with Unknown HIV status. On the other hand variables like Age and type of tuberculosis were not found as predictor factors.

In Zigong, women in the age category of 15-49 years have 1.55 more likely to show treatment success (95% CI: 1.08, 2.206) compared with older age. But, women with MDR TB were less likely to show treatment success (Adjusted OR, 0.90; 95%CI: 0.24, 0.34) compared with non-drug resistance cases. (Table: 4)

**Table 4.** Multiple Logistic regression factors affecting treatment outcome of women in Tigray and Zigong January 2007 December 2016 N=2084 and N= 4047.

| Tigray |  |  |  |  |  | Zigong |  |  |  |  |
| --- | --- | --- | --- | --- | --- | --- | --- | --- | --- | --- |
| Characteristic | Total | Un success<br>N% | COR(95%CI | AOR(95% CI) | P-value | Total | Un success<br>N% | COR(95%CI) | AOR(95% CI) | P-value |
| **Type of TB** |  |  |  |  |  |  |  |  |  |  |
| P-Positive | 528 | 51(9.6) | 0.55(0.38,0.81)* | 0.89(0.58,1.37) | 0.61 | 1889 | 60(3.2) | 1.023(0.72,1.45) | 0.91(0.63,1.31) | 0.614 |
| P-Negative | 1062 | 83(7.8) | 0.7(0.50,0.97)* | 0.78(0.56,1.09) | 0.15 | 2156 | 70(3.2) | 0.03(0.002,0.54) | 0.025(0.002,0.41) | 0.01 |
| E. pulmonary | 1214 | 71(5.8) | 1 | 1 |  | 2 | 1(50) | 1 | 1 |  |
| **T. Category** |  |  |  |  |  |  |  |  |  |  |
| Re-treatment | 95 | 24(25.2) | 0.28(0.13,0.34)* | 0.29(0.16-0.53) | 0.0001 | 3822 | 122(1.7) | 0.82(0.41,1.62) | 1.12(0.53,2.35) | 0.76 |
| New | 2709 | 178(6.5) | 1 | 1 |  | 225 | 9(4.0) | 1 | 1 |  |
| **Age** |  |  |  |  |  |  |  |  |  |  |
| 15-49 | 2155 | 156(7.2) | 0.98(0.69,1.37) | 1.17(0.81,1.69) | 0.378 | 2494 | 69(2.8) | 1.46(1.03,2.07) | 1.55(1.08,2.206) | 0.016 |
| ≥ 50 | 649 | 46(6.9) | 1 | 1 |  | 1553 | 62(4.1) | 1 | 1 |  |
| **Drug resistance** |  |  |  |  |  |  |  |  |  |  |
| MDR | 7 | 6(75) | 0.013(0.002,0.11) | 0.31(0.003,0.27) | 0.002 | 13 | 3(31.2) | 0.109(0.03, 0.42) | 0.090 (0.024,0.34) | 0.0001 |
| NMDR | 2797 | 196(7.0) | 1 | 1 |  | 4034 | 128(3.2) | 1 | 1 |  |
| **HIV status** |  |  |  |  |  |  |  |  |  |  |
| Negative | 1838 | 88(4.7) | 3.39(2.37,4.87)* | 3.58(2.47,5.18) | 0.0001 | ND | ND | ND | ND |  |
| Positive | 596 | 60(10) | 1.53(1.03,2.26)* | 1.59(1.07,2.38) | 0.022 | ND | ND | ND | ND |  |
| Not tested | 370 | 54(14.5) | 1 | 1 |  | ND | ND | ND | ND |  |
| **Total** | 2804 | 202 |  |  |  | 4047 | 131 |  |  |  |

### Findings and Discussion

Tuberculosis is exacerbated by malnutrition and frequently affects economically active young adults (24, 25). Thus, women of reproductive age are more likely to develop active TB if they encounter TB bacteria and they are less likely to seek help for TB symptoms than men (4). In this study the mean age of TB case in Tigray was 36 years with SD±15 and in Zigong was 44years with SD±17. Tigray finding was similar with study done in sidama and Gojjam for general population (17, 20). And comparing with Zigong, younger women get infected in Tigray than Zigong and this could be the age distributions tuberculosis in Africa has been severely skewed by the human immunodeficiency virus epidemic (26). Besides, under nutrition was a major public health problems in Tigray and it has considerable effect in provoking tuberculosis infection (27, 28). The demographic structural difference may be other cause large number of old peoples were found in Zigong than Tigray this all reasons may be factors to have more young age TB cases in Tigray than Zigong.

In 2014, globally TB killed 480,000 women and 140,000 children (2). Correspondingly, in this retrospective study the death of women and children in the past ten years were 170 [6 %] in Tigray and 36 [0.8%] in Zigong. The death toll in Tigray was higher than the annual report of the Regional Health Bureau for both sex 3.6% (33), retrospective study done in Gojjam for both sex 3.7% and global report of TB death for women and children 4.2% (10, 14, 17). In cases of Zigong it is similar with the global report for general population (10). Comparing with Zigong, more death occurred in Tigray. This could be the prevalence of TB/HIV co morbidity was high in sub-Saharan country(34) or poor women health care seeking behavior and diagnostic delay (35) Poor quality of treatment in High TB burden countries(36) are in favor of bad treatment outcome.

Treatment success is sum of patients cured and those who have completed treatment. Hence, patient compliance is a key factor in treatment success (10). In this study, the overall treatment success rate of all TB cases was 92% in Tigray and 96.6% in Zigong. The finding of Tigray was similar with studies done in west Gojjam, Sidama and Addis Ababa 91.5% (17, 20, 37) the success rate for both sex. But, higher than the WHO report 89% for both sex (10) and the success of Zigong is higher than (10, 38, 39). Thus, the better treatment success in both countries could be since these studies assess only women and this group have good treatment adherence as a result the percentage of treatment success was better comparing with general population.

A patient is considered “cured” when sputum smear examination is bacteriologically negative in the last month of treatment and on at least one previous occasion(17). The cure rate for pulmonary positive cases was 90% in Tigray and 95% in Zigong. So, the finding of Tigray was slightly higher than 2015/16 annual report of the region and study done in west Gojjam (14, 17, 33) for general population. But, in case of Zigong it is similar with the WHO global report for Chinese population (10). Therefore, the reason of higher cure rate in Zigong could be the less number of HIV co infected women and/or the early initiation of drug resistance test program in the region were made Zigong to have higher cure rate than Tigray.

Retreatment case is patient who has been treated for one month or more with anti-TB drugs in the past(2) and many studies indicate that re treatment case have high chance of poor treatment outcome or failure rate. Similarly, in our study re-treatment cases were 79% less likely (adjusted OR, 0.29; 95% CI: 0.16-0.53) to have treatment success compared to new cases and this finding was parallel with the study done in Gojjam, Sidama, Tigray and Uganda (13, 17, 20, 40). Yet, in Zigong it was not significant this may be the number of re treatment cases were few Zigong compared with Tigray. Besides, drug sensitivity test is done in Zigong before providing anti-TB drug to a patient and better follow up.

Globally, an estimated 3.3% of new TB cases and 20% of previously treated cases have MDR-TB (2). In Ethiopia 2.7% of the new and 14% of the previously treated TB cases expected to have had rifampicin or multi drug resistant TB (10). Report about MDR-TB started in 2011 in both countries. Then in Tigray total of 28(0.4%) MDR-TB cases were reported and majorities were identified in 2016 which is 16(3.7%) cases. Where as in Zigong 16 (0.3%) MDR-TB cases were identified. According to global TB report only 50% of MDR-TB patients were successfully treated(10). But in this study 6 women out of 7 MDR TB cases in Tigray and 3 women out of 13 MDR TB cases in Zigong were not successfully treated. this is consistent with study done in India China and Ethiopia (41, 42). In both study area patients with drug resistant tuberculosis were less likely to have treatment success. In Tigray 69% less-likely (Adjusted OR, 0.31; 95% CI: 0.003, 0.27 and in Zigong 9% less likely (adjusted OR, 0.09; 95% CI: 0.024, 0.34 successful treatment outcome compared with non-drug resistant. It is obvious that the nature of the bacteria, the long-term treatment and its adverse effect made the outcome poor and many studies support this finding (25, 43).

## Conclusions

Evidence presented in this study shows tuberculosis is one of the major public health concerns for women in both countries. However, poor level of treatment success and mortality were seen in Tigray compared with Zigong. Besides, factors boost unsuccessful treatment outcome were many in case of Tigray than in Zigong. Hence, national policy makers of Ethiopia and region Tigray should give due attention to this specific group.

## Abbreviation

TB: Tuberculosis
HIV: Human immune deficiency Virus
MDR/TB: Multi drug resistance tuberculosis
CDC: The Centers for Disease Control and Prevention
WHO: The World Health Organization

## Authors’ contributions

The authors’ contribution was as described below. XM and MG conceived and designed the study. MG and BA performed the study. MG, XM, and BA analyzed the data. MG XM and MF wrote the manuscript. All authors read and approved the final manuscript.

## Acknowledgments

The authors would like to thank Sichuan University and Mekelle University for funding this study. We also would like to thank Zigong CDC, Tigray Regional Health Bureau and TB treatment centers of each country providing the data. We would like also to extend our thanks to all the data collectors

## Author details

1 Department of Health-Related Social and Behavioral Science, West China School of Public Health, Sichuan University, Chengdu 610041, China; 2 School of Public Health, college of Health Science Mekelle, University, Tigray, Ethiopia

